# Ex vivo massively multidimensional diffusion-relaxation correlation MRI: scan-rescan reproducibility and caveats

**DOI:** 10.1101/2025.03.17.643705

**Authors:** Pak Shing Kenneth Or, Maxime Yon, Omar Narvaez, Eppu Manninen, Tarja Malm, Alejandra Sierra, Daniel Topgaard, Dan Benjamini

## Abstract

Massively multidimensional diffusion-relaxation correlation MRI (MMD-MRI) provides information beyond the traditional voxel-averaged metric that may better characterize the microstructural characteristics of biological tissues. MMD-MRI reproducibility has been established in clinical settings, but has yet to be thoroughly evaluated under preclinical conditions, where superior hardware and modulated gradient waveforms enhance its performance. In this study, we investigate the reproducibility of MMD-MRI on a micro-imaging system using ex vivo mouse brains. Notably, the estimated signal fractions of intra-voxel spectral components in the MD-MRI distribution, corresponding to white and gray matter, along with the frequency-dependent parameters, demonstrated high reproducibility. We identified bias between scan and rescan in some of the metrics, which we attribute to the time gap between repeated scans pointing to a long-time progressive fixation effect. We compare our results with in vivo results from clinical scanners and show the reproducibility of diffusion frequency-dependent metrics to benefit from the improved gradient hardware on our preclinical setup. Our results inform future micro-imaging ex vivo MMD-MRI studies of the reproducibility of MMD-MRI metrics and their dependence on fixation time.

## 1. Introduction

In medical magnetic resonance imaging (MRI), tissue microstructure imaging is the characterization of tissue at the micrometer scale, which is typically two-to-three orders of magnitude finer than in vivo clinical MRI resolution^1^. The feat is challenging because of the complexity of tissue, which can consist of multiple, interacting or non-interacting spin populations^2,3^ which behaviors have not been fully characterized for different tissue types. This can be addressed through acquisition schemes that combine various diffusion encoding magnitudes and shapes^4^, quantitative relaxometry^5^, and tissue-specific modeling^6-8^. Across different areas of nuclear magnetic resonance (NMR) and MRI applications, it has been shown that acquisition schemes that correlate two or more MR parameters can yield information that would otherwise remain inaccessible if the parameters were measured independently^9-12^. A multidimensional MR measurement also enables the accounting of inherent biases that the combining one-dimensional measurements would have produced had they been acquired independently^5^. For example, signal fractions from different spin populations obtained by inversion using any model that does not account for relaxation may be mis-estimated in edema-related conditions^7^. The wealth of information obtainable from diffusion-relaxation correlation studies has been demonstrated in detecting diffuse axonal injury^13^, spinal cord injury^14,15^, neuroinflammation^16^, placenta dysfunction^17^, breast^18^ and brain cancer^19^, thus providing this modality great potential for patient diagnosis.

Another MRI modality that has been a topic of interest is diffusion spectrum encoding^20^. It is a method used to characterize the time dependence of diffusion behavior, which arises when water molecules diffuse in the presence of barriers that restrict their motion as is likely the case in complex biological tissue. By varying the diffusion encoding waveform modulation or duration, the probed spectra of diffusion times, or diffusion frequencies, is also varied to control the amount of interaction between the water molecules and the barriers. Information about the barriers can then be extracted from the time dependence, adding an extra dimension of tissue characterization. Its utility has been demonstrated in understanding the mechanism of acute ischemic stroke^21^, understanding epidermoid cysts^22^, grading of intra-axial brain tumors^23^ and more.

The interpretation of a measurement can either be done through fitting to a biophysical model with assumptions specific to the measured tissue, or a general approach that makes less assumptions. While the biophysical modelling approach provides great interpretability to the results, incorrect model assumptions will lead to mischaracterization and misinterpretation^7,24-26^. A general non-parametric approach does not suffer from this pitfall, but is computationally expensive and prohibits direct use of a forward signal model for various purposes such as protocol optimization^27^. Nevertheless, for applications to samples that are as complex and heterogeneous as, for example, different types of tissue across the human brain, the cost of a non-parametric approach can be a small price to pay to mitigate the risk of inaccurate model assumptions.

Massively multidimensional diffusion-relaxation correlation MRI (MMD-MRI)^28^ is a technique that integrates several above-mentioned MRI approaches into a single protocol. Tensor-valued diffusion encoding^29^ is combined with diffusion spectrum encoding to probe the distribution of frequency-dependent diffusion tensors^30^, which are further correlated with relaxation measurements^28^, while using a general parameter estimation method (Monte Carlo inversion^31,32^) to untangle the multiple dimensions. As an all-in-one approach, we expect the technique to be able to characterize a wide range of tissue types and be useful in many research scenarios.

An efficient and sparse in vivo MMD-MRI clinical acquisition protocol that provides whole brain coverage at 2-mm isotropic resolution was recently introduced^33^. The reliability and repeatability of the clinical MMD-MRI framework has been established, showing it to be comparable with conventional diffusion MRI reproducibility^34^. Human studies are essential for advancing the translation and application of MMD-MRI in clinical practice but have inherent limitations compared to preclinical animal models. Preclinical neuroimaging offers distinct advantages, including controlled experimental conditions, the ability to use complementary invasive techniques, access to genetic and disease models, and superior imaging hardware. These benefits are particularly valuable for studying the mechanisms underlying brain function and pathology. Recently, the application of MMD-MRI on a preclinical setup has been demonstrated on rats ex vivo^28^ and in vivo^28,35^. Before applying this technique to systematic comparative studies, our aim in the current study is to establish the reproducibility of a preclinical MMD-MRI framework by characterizing ex vivo mouse brains and assessing intra-scanner scan-rescan reproducibility.

## 2. Methods

### 2.1 Samples

Eight female mice, four of which were wild type and four were 5xFAD (a transgenic model to study familial Alzheimer’s disease^36^), were perfused at approximately 8 months old with 0.9% saline for 2 min at 5 ml/min followed by phosphate-buffered 4% paraformaldehyde solution for 20 min at 5 ml/min. The brains were extracted, trimmed to fit in 10 mm NMR tubes, preserved in phosphate-buffered 2% paraformaldehyde solution and stored at 5°C. The animal procedures were approved by the Animal Ethics Committee of the Province Government of Southern Finland and carried out according to the guidelines set by the European Community Council Directives 2010/63/EEC.

### 2.2 MRI acquisition

All scans were acquired on a Bruker Avance-III 11.7 T spectrometer (equipped with a MIC-5 probe, 3 T/m maximum gradient amplitude) with the preclinical imaging software ParaVision 360. For each sample, two MMD-MRI acquisitions with EPI readout, partial Fourier factor 6/8, and 3D k-space encoding of the same protocol were acquired at 150-µm isotropic resolution. The scans were two to seven months apart. Among other parameters, the MMD-MRI protocol acquires *b* values from 0 to 8•10^9^ sm^-2^, echo time τ_E_ 9.4 to 49.4 ms, and repetition time (time between saturation pulses after one acquisition and excitation pulse of next acquisition) τ_R_ from 0.8 to 3.5 s (Figure 1). As was the case in previous preclinical MMD-MRI studies^28,37^, strong gradients enabled the use of modulated diffusion gradients^38^ that have high and narrow ranges in their encoding spectra while still achieving strong enough *b* values within acceptable τ_E_s. Following the convention of Jiang et. al.^38^, waveforms with modulation orders 0 to 2 were used in this study.

**Figure 1.**
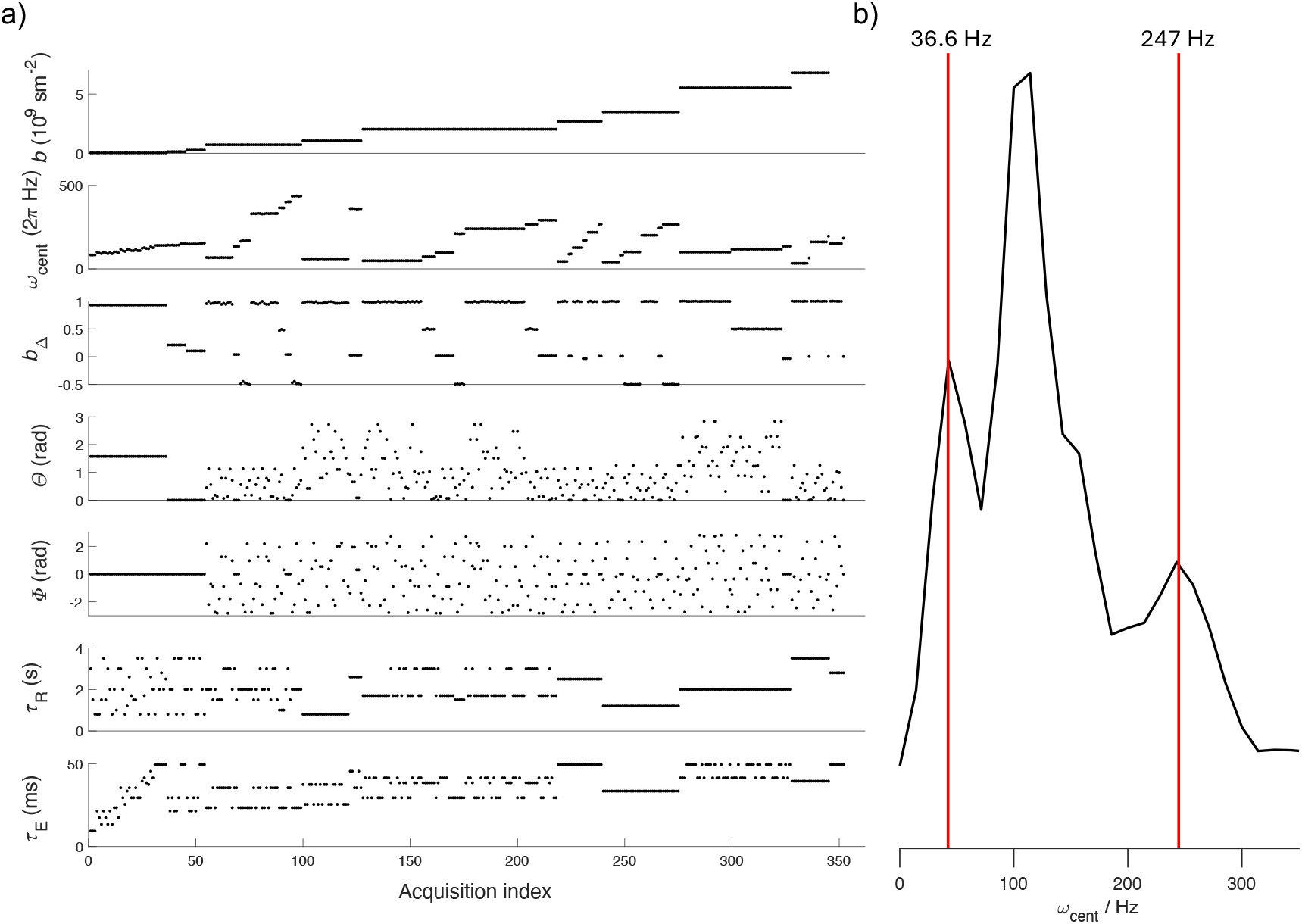
(a) MMD-MRI acquisition protocol of 352 images, acquired at different combinations of *b*-values, centroid frequencies (*ω*_cent_) ^38^, diffusion tensor shapes (*b*_Δ_), orientations of the diffusion encoding tensors (*Θ, Φ*), repetitions times (*τ*_R_) and echo times (*τ*_E_). (b) *b*-value-weighted distribution of *ω*_cent_ of the protocol, effectively excluding low *b*-value contributions. Red lines depict the 10^th^ percentile and the 90^th^ percentile.

At each parameter setting, two scans with reversed phase encoding directions were acquired to correct for EPI distortion. While both scans were done at the same temperature setting (298 K), the flow rate of the variable-temperature (vt) gas (which controls the sample temperature) was changed from 400 Lph for the first scans to 200 Lph for the second scans to reduce motion of the sample during acquisition. No effect on the data was observed.

Additional to the MMD-MRI acquisitions, a standard 3D FLASH scan at 50 µm resolution was acquired per sample with τ_E_ = 10 ms, repetition time (time between excitation pulse of one acquisition and excitation pulse of the following acquisition) 0.7 s and flip angle = 30° to facilitate image registration.

### 2.3 Data processing

#### 2.3.1 Preprocessing

After image reconstruction on ParaVision 360, denoising was done with MP-PCA^39^ with a patch size of 7×7×7, Gibbs ringing removal was done on MRtrix3^40^, and EPI distortion correction was done using the TopUp function from FSL^41^.

The FLASH acquisitions were not preprocessed.

#### 2.3.2 Parameter estimation: MMD-MRI inversion

As is done in other MMD-MRI studies^28,34,35^, the estimation of MMD-MRI parameters^30^ is performed using the multidimensional diffusion MRI toolbox for MATLAB^42^, which employs a Monte Carlo inversion strategy^31,32^ to solve the inverse problem. The toolbox assumes the MMD-MRI acquisition signal *S* in each voxel is represented by a sum of mono-exponential components *i*

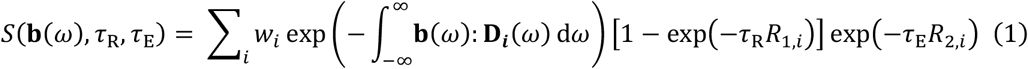

where ***b***(*ω*) is the tensor-valued encoding spectrum, *w*_*i*_ the weight of each component, **D**_*i*_(*ω*) the tensor-valued diffusion spectra^43^ in the laboratory frame of reference, and *R*_*1,i*_ and *R*_2,*i*_ the longitudinal and transverse relaxation rates of each component respectively. Each component is assumed to have axial symmetry in its principal axis system, which is rotated into the lab frame of reference through a rotational transformation **R**(*θ*_*i*_, *ϕ*_*i*_),

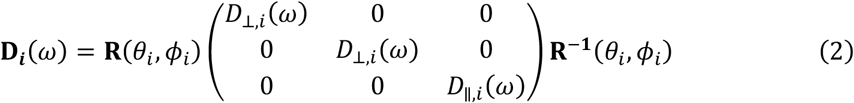

The function forms of *D*_⊥,*i*_(*ω*) and *D*_∥,*i*_(*ω*) are parametrized with Lorentz distributions, characterized by the high frequency (i.e. restriction-free diffusion) isotropic diffusivity *D*_0,*i*_ and, axial and radial transition frequencies (Γ_∥,*i*_ and Γ_⊥,*i*_)^30,43^ as

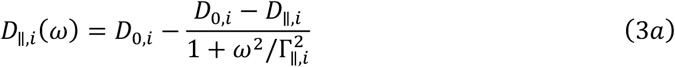

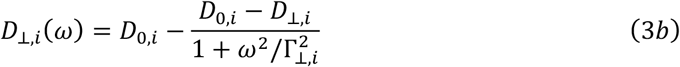

For convenience of interpretation, the diffusivity parameters are expressed as isotropic diffusivity *D*_iso,*i*_(*ω*) and normalized diffusion anisotropy *D*_Δ,*i*_(*ω*) by transforming *D*_⊥,*i*_(*ω*) and *D*_∥,*i*_(*ω*)

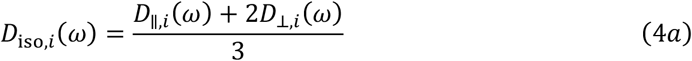

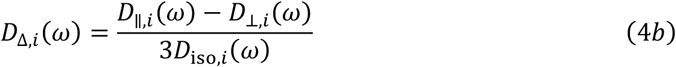

Employing a bootstrapping strategy, an ensemble of solutions in the **D**(*ω*) − *R*_1_−*R*_2_ space is obtained for the same measurements. The ensemble of solutions collectively assesses the uncertainty in the solution landscape^31^. The solutions can be put together to form a continuous distribution of components^31,32^, which can further be quantified in terms of means (E[*x*]), variances (V[*x*]), and covariances (C[*x, y*]) at any specific *ω*. The analysis of the distributions is also done in bins in the 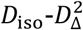 space^44^, which are defined as follows: bin 1: 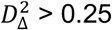 and 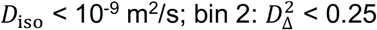 and *D*_iso_ < 10^−9^ m^2^/s; bin 3: *D*_iso_ > 10^−9^ m^2^/s. Within the brain, these boundaries separate white matter (WM) from gray matter (GM), and the two from freely diffusing cerebrospinal fluid.

To evaluate how strongly *D*_iso_(*ω*) and *D*_Δ_(*ω*) depend on *ω*, their E[*x*], V[*x*] and C[*x, y*] values at two different frequencies (*ω*_*min*_ and *ω*_*max*_) are evaluated to compute Δ_*ω/ 2π*_ metrics

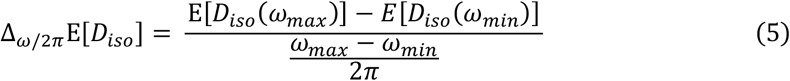

and similarly, for V[*x*] and C[*x, y*] metrics, and for 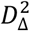.

Standard diffusion-encoded color (DEC) maps^45^ were made by finding the “median” fractional anisotropy (FA) and the “median” eigenvector. For each bootstrap solution, an FA and an eigenvector is defined by its mean diffusion tensor 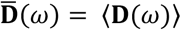,so a “median” can be defined over the bootstrap solutions. Notice that the mean diffusion tensor is a function of *ω*, so FA is also a function of *ω*0

In this approach, the only settings are the inversion limits of the solution space and inversion algorithm settings. For our data, the inversion limits used are 5×10^−12^ m^2^s^-1^ < *D*_0*/*⊥*/*∥_ < 5×10^−9^, 0.1 s^-1^ < Γ_⊥*/*∥_ < 10^5^ s^-1^, 0.1 s^-1^ < *R*_1_ < 4 s^-1^, 4 s^-1^ < *R*_2_ < 100 s^-1^. The other inversion algorithm settings are identical to that in Narvaez et al.^28^. Approximately, the inversion time for 130000 voxels on a system with two 24-core, 2.65 GHz CPUs was 40 hours.

#### 2.3.3 Consolidation of bootstrap solutions and ROI-averaging

Two-dimensional projections of the full **D**(*ω*)-*R*_1_-*R*_2_ distributions were constructed to facilitate visualization and to allow investigation of certain regions of interest (ROIs). To obtain the distributions in each voxel, the bootstrap solutions first have to be consolidated into a single representative distribution. This was done using the following procedure: (1) grouping **D**(*ω*)-*R*_1_-*R*_2_ components across different bootstrap solutions using *k*-means clustering over 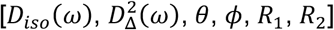 as features (after normalization according to their respective maximal values) with the *L*1 distance metric. (2) Once grouped, each cluster is transformed into a single **D**(*ω*)-*R*_1_-*R*_2_ component by taking the median of each parameter (e.g., *D*_iso_), and the median of the weights. This process then allowed the projection and mapping of the median weights of the discrete components from the consolidated distributions onto 64 × 64 meshes in the 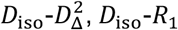,and *D*_iso_-*R*_2_ planes.

These voxel-wise 2D projections were then summed over entire ROIs within each scan, normalized, and finally averaged across all scans to produce characteristic distributions.

#### 2.3.4 Image registration and Region of interest (ROI) definition

By registering all scans to a mouse brain atlas, ROIs in native space were obtained for analysis, as seen in Figure 2 and detailed in Table 1. The atlas used is the Turone Mouse Brain Template and Atlas (NITRC – www.nitrc.org). The atlas provides a reference structural scan, which was used for registration, and labels that delineate brain regions on the structural scan.

**Table 1.**
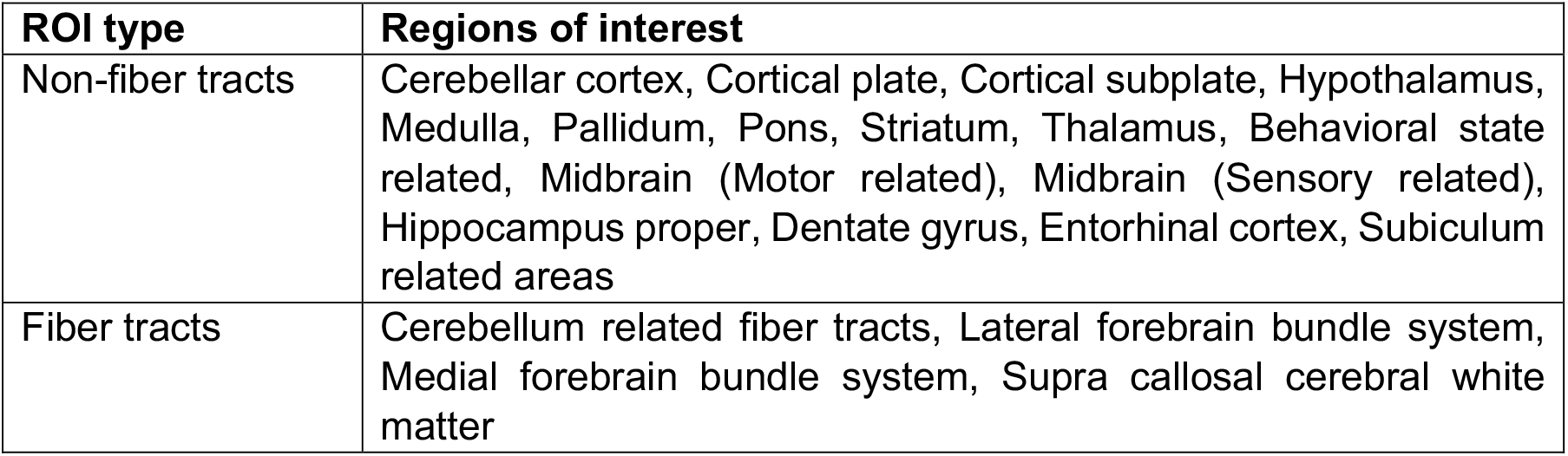
Regions of interest (ROIs) delineated in data with atlas. While finer ROIs are available from the atlas, they are impractical to use due to the difference in resolution between it and the MMD-MRI scans (60 µm vs 150 µm).

| ROI type | Regions of interest |
| --- | --- |
| Non-fiber tracts | Cerebellar cortex, Cortical plate, Cortical subplate, Hypothalamus, Medulla, Pallidum, Pons, Striatum, Thalamus, Behavioral state related, Midbrain (Motor related), Midbrain (Sensory related), Hippocampus proper, Dentate gyrus, Entorhinal cortex, Subiculum related areas |
| Fiber tracts | Cerebellum related fiber tracts, Lateral forebrain bundle system, Medial forebrain bundle system, Supra callosal cerebral white matter |

**Figure 2.**
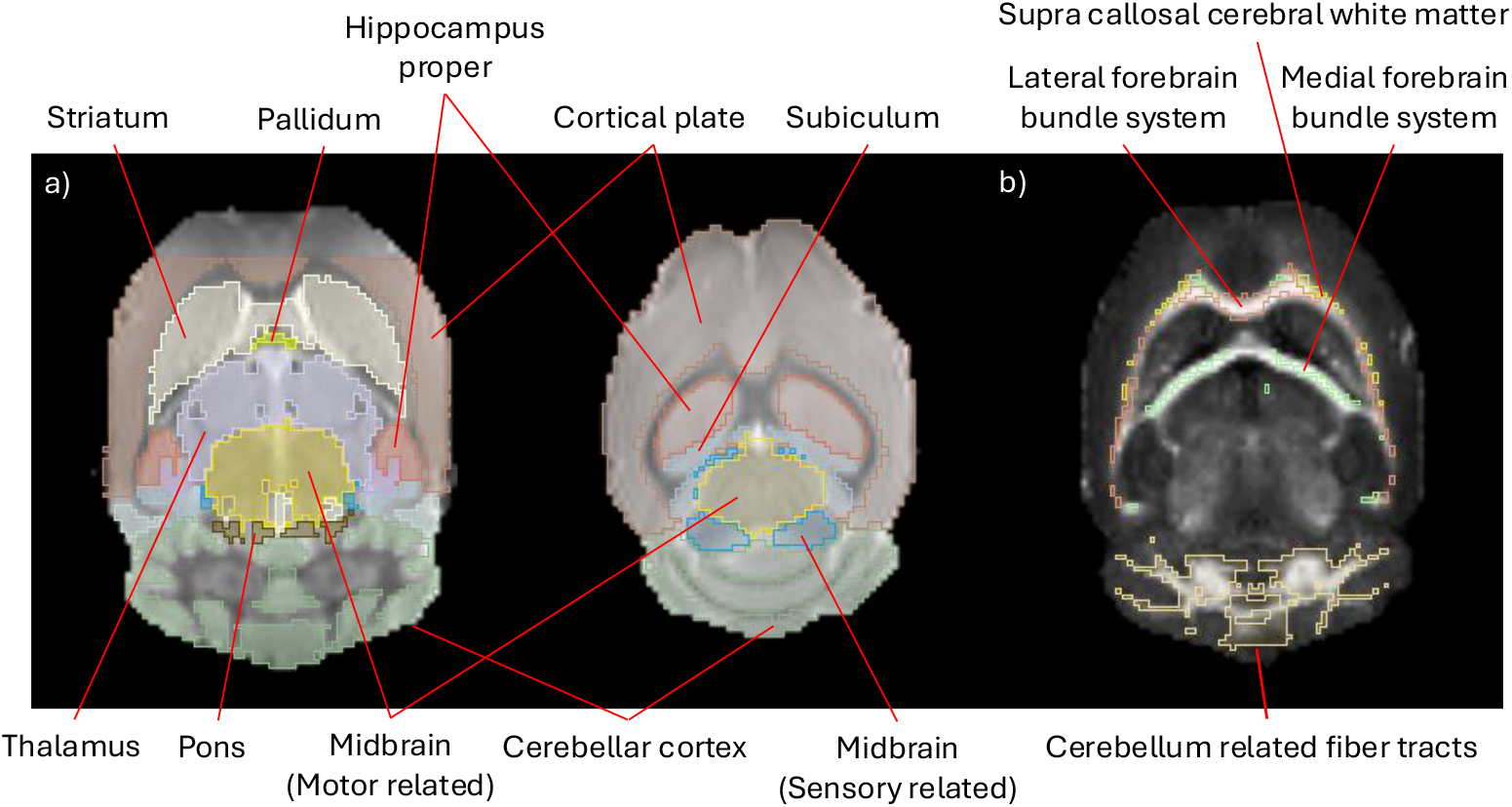
(a) Gray matter ROIs overlayed on top of a b0 structural image. (b) White matter fiber ROIs overlayed on top of a 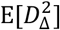 image.

Advanced Normalization Tools^46^ (ANTs version 2.5.1, http://stnava.github.io/ANTs/, accessed on 22 April 2024) was used for template creation and image registration. In brief, registration to the atlas reference consists of multiple steps, including creation of a mid-way template between scan and rescan, and a study-specific FLASH template. This was done utilizing three scans/maps, including a ‘b0’ structural image for each scan, a DEC map for each scan, and a high-resolution FLASH scan for each sample. The ‘b0’ structural image was made by summing the images from six scans within the MMD-MRI protocol where *b*-value = 0 such that a structural image representing that scan is obtained. Inclusion of the DEC maps enabled better registration of thin WM fibers. See supplementary materials for a detailed description of the registration steps.

Regions of interest were grouped by brain region as defined by the Turone atlas (Table 1). The exceptions for this are the hippocampus proper, dentate gyrus, entorhinal cortex and subiculum, which belong to the hippocampal formation.

#### Reproducibility analysis

The reproducibility of each metric was assessed by comparing the ROI-averaged value of the metric from the first scan of one sample to the repeat scan of the same sample. For binned metrics, the bin fraction-weighted average is taken instead. Correlation plots were made for a qualitative assessment, and Lin’s Concordance Correlation Coefficient^47^ (CCC) was computed for a quantitative assessment. CCC is a typical statistic used to measure reproducibility, taking not only Pearson’s correlation into account but also the change in mean and variance between scan and rescan in its evaluation. The advantage CCC has over Pearson’s correlation coefficient is that a high CCC can be achieved not only when the repeated measurements lie on a straight line, but also the measurements have to be the same^48^.

Bland-Altman plots were created to investigate the presence of bias by using MATLAB class “Bland-Altman and Correlation Plot”^49^. Because the differences between scan and rescan were not Gaussian, the limits of agreement (LOA) were estimated with the sample quantile estimator^50^.

To be elaborated in the Discussion section, a systematic shift was observed in some of the metrics. The shifts were sample-dependent and not related to the acquisition or processing. To factor out the bias, for each metric, Pearson’s correlation coefficient was calculated for each sample to make boxplots for assessing “method-specific” reproducibility.

To facilitate the comparison of reproducibility with other studies, the within-subject coefficient of variation is computed. Following the definition in Manninen et al.^34^, *CV*_ws_ is estimated by

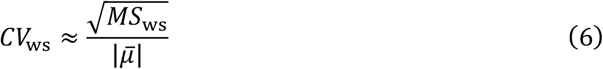

where *MS*_ws_ is the within-subject mean square difference and 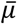 is the mean over samples and scans.

## 3. Results

### 3.1 Regional microstructural characterization of the mouse brain

Figure 3 shows projections of the bootstrap solution distribution in **D(**ω)-*R*_1_-*R*_2_ space onto two and one dimension(s), obtained from single voxels and from entire ROIs. Distributions of the entire ROIs are obtained according to the procedure described in the Methods section. Projections from three selected regions are presented to highlight their distinct characteristics.

**Figure 3.**
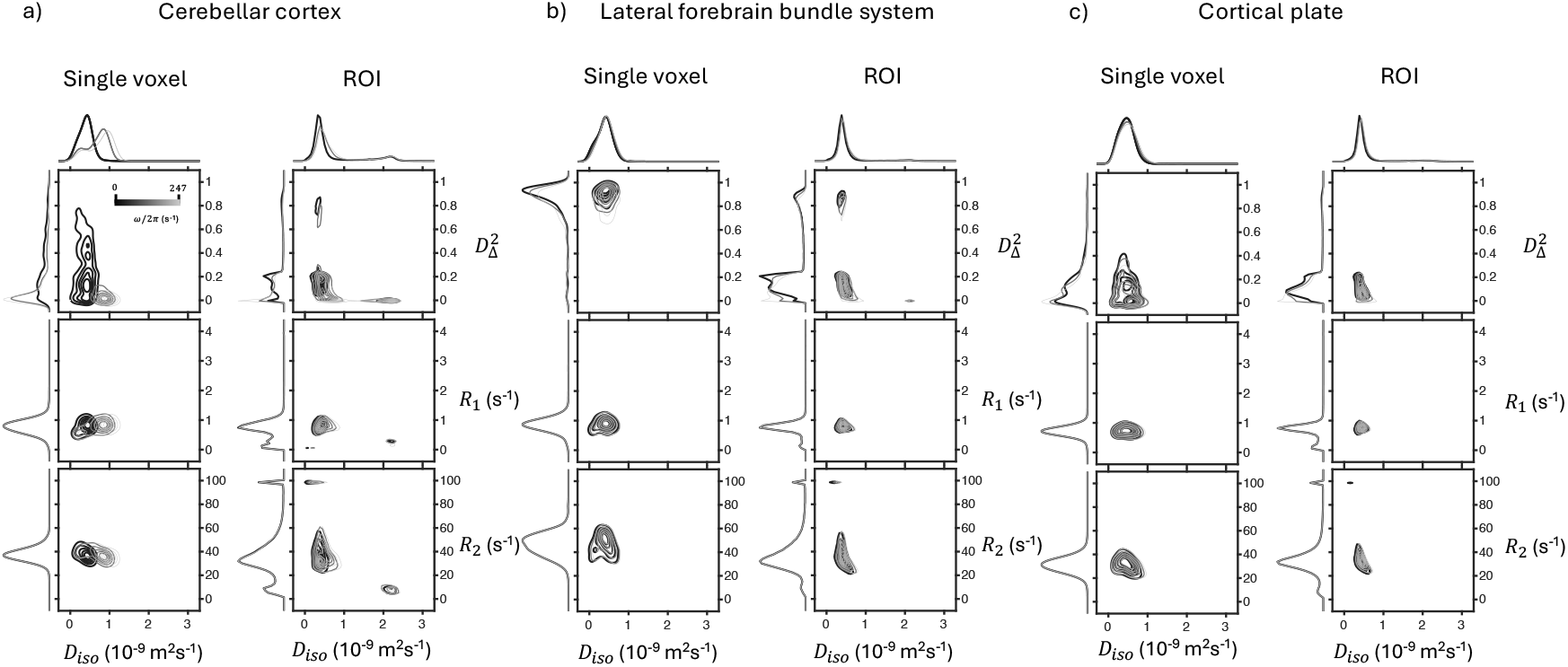
2D projections on to the 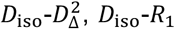and *D*_iso_-*R*_1_ dimensions of the complete the solution distribution in **D**(*ω*)-*R*,-*R*_1_ space, from a single voxel (left column of each sub-figure) and from an entire ROI (right column of each sub-figure). On the edges of each plot are 1D projections to the corresponding dimension. Distributions at different diffusion frequencies *ω/*2*π* are plot with different shades of black. Here the distributions from the (a) cerebellar cortex, (b) lateral forebrain bundle system and (c) cortical plate are shown.

The cerebellar cortex is shown for its strong *D*_iso_ and 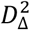 frequency dependencies (i.e., 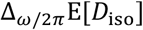 and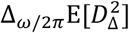). This can be seen in the 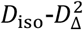 projection panels most easily in the single voxel distribution (Figure 3a, left column). The 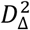 changes from a wide distribution between 0 and 0.6 at the highest frequency, to a singular peak near 0 at the lowest frequency. Similarly, *D*_*iso*_ splits from a singular peak at the lowest frequency to two at the highest frequency. The lateral forebrain bundle system is shown as an example of highly anisotropic microstructure, illustrated in the single voxel 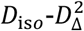 panel, the distribution is centered near the maximum value of 1 (Figure 3b, left column). The cortical plate was shown as an example of gray matter, which is characterized by low 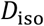 and 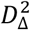,and a moderate response to diffusion encoding frequency. Two fractions, separable by 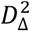,are detected, which merge at higher frequencies.

On the ROI-averaged projections (Figure 3, right columns), we see a narrowing of peaks on the 1D projections which reflects the typical improvement of SNR by taking a larger sample size. However, ROI-averaged results are also affected by partial volume effects from voxels at the boundaries of the regions, which possibly result in extra, highly separate fractions that are not seen in the single-voxel results. This is most severe in the lateral forebrain bundle system ROI, due to its long and narrow nature, creating a low 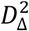 fraction that is much larger than the high 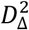 peak. The small, high *R*_1_ and high *R*_2_ peaks are inversion artefacts from a small number of voxels that mostly lie at the boundary between sample and the surrounding media).

The voxelwise distributions can be summarized by their means (E[*x*]), variances (V[*x*]), and covariances (C[*x, y*]) over the entire solution space. This straightforward dimensionality reduction allows for visualization of the essential relaxation-diffusion characteristics in a representative slice from one of the ex vivo mouse brains, as shown in Figure 4.

**Figure 4.**
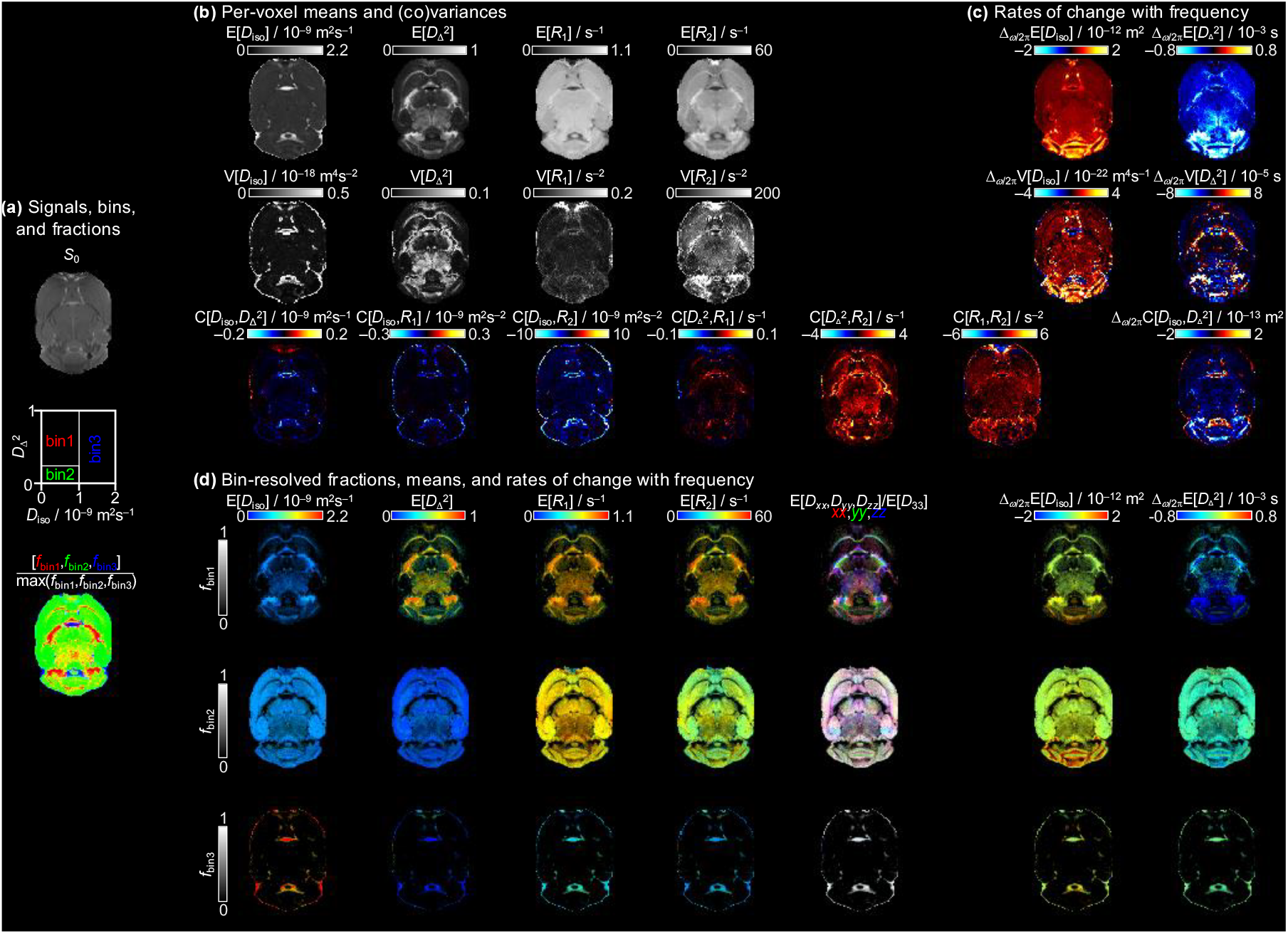
Parameter maps in a transvers slice of a scan. (a) Signal amplitude map estimated by the solution, binning scheme and dominant bin composition of each voxel. (b) Per-voxel means, variances and covariances. (c) Diffusion encoding frequency dependence of *D*_iso_ and 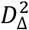,as described by equation (3). (d) Bin-resolved, per-voxel means of MRI metrics.

### 3.2 Test-retest reproducibility

Figure 5 shows the correlation plots and CCCs for the investigated parameters. To enhance readability, the plot layout aligns with the parametric maps in Figure 4. In general, plots that have most of their data points near the line of equivalence have high CCCs. Following the recommendation of McBride^48^, we rate the reproducibility of each parameter according to their CCCs, summarized in Table 2.

**Table 2.** MMD-MRI metrics rated by reproducibility based on CCC. Poorly performing metrics (CCC < 0.900) are not listed.

| Strength-of-agreement | Metrics |
| --- | --- |
| Almost perfect (>0.99) | $\Delta_{\omega/2\pi} E[D_{iso}]_{bin2}$ |
| Substantial (0.95-0.99) | $E[D_{\Delta}^2]$, $E[R_2]$, $V[D_{\Delta}^2]$, $\Delta_{\omega/2\pi} E[D_{iso}]$, $\Delta_{\omega/2\pi} E[D_{\Delta}^2]$<br>$Fraction_{bin1}$, $E[D_{\Delta}^2]_{bin1}$, $\Delta_{\omega/2\pi} E[D_{iso}]_{bin1}$, $\Delta_{\omega/2\pi} E[D_{\Delta}^2]_{bin1}$<br>$Fraction_{bin2}$, $E[D_{\Delta}^2]_{bin2}$ |
| Moderate (0.90-0.95) | $E[D_{iso}]$, $V[D_{iso}]$, $\Delta_{\omega/2\pi} V[D_{iso}]$, $\Delta_{\omega/2\pi} V[D_{\Delta}^2]$, $\Delta_{\omega/2\pi} C[D_{iso}, D_{\Delta}^2]$,<br>$E[R_2]_{bin1}$, $E[R_2]_{bin2}$ |

**Figure 5.**
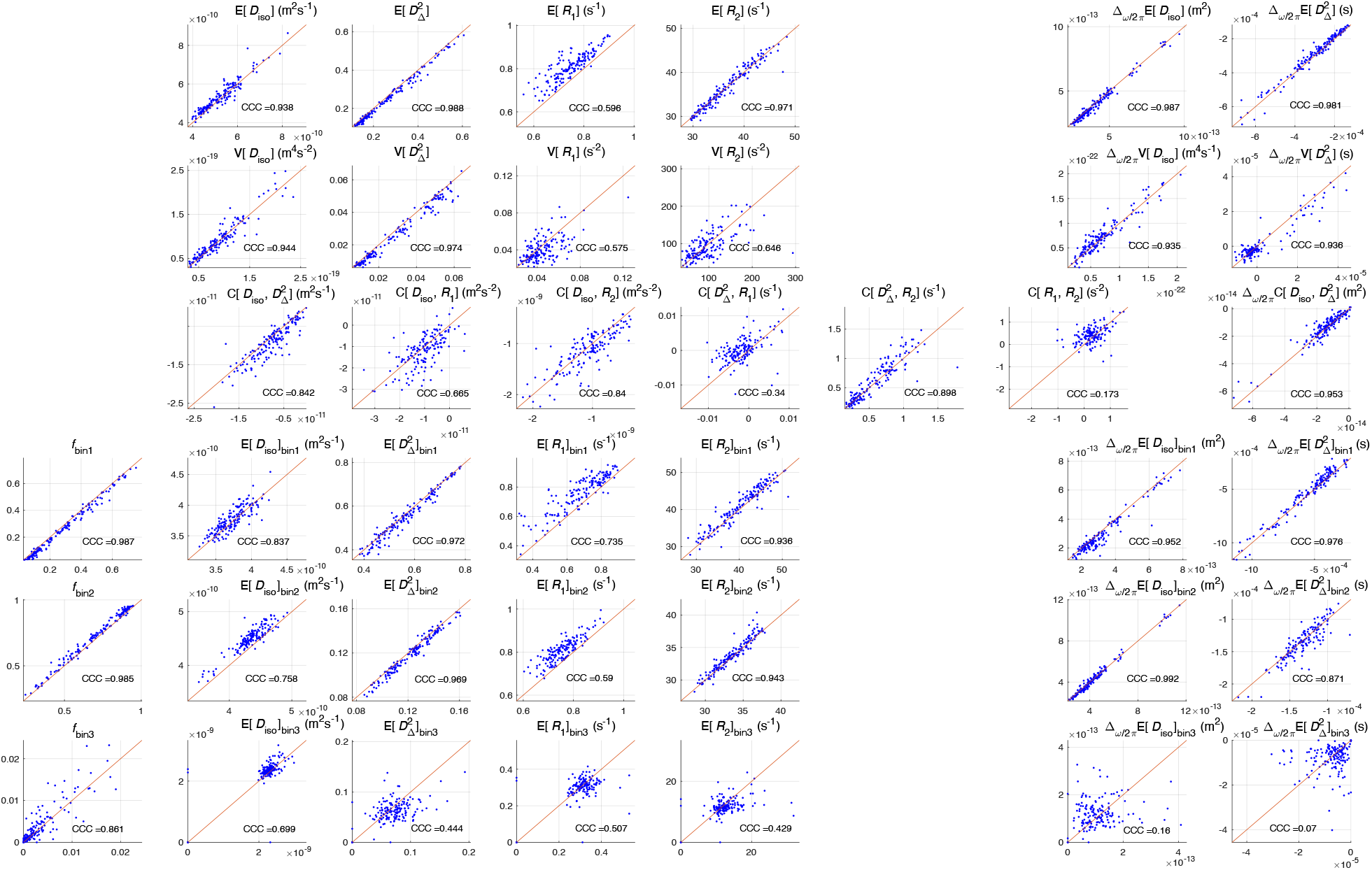
Correlation plots of MMD-MRI metrics and their CCCs for per-voxel metrics and bin-resolved per-voxel metrics, following the layout of Figure 4. Each point on the plot is the ROI-averaged value of the metric from a sample from one of the ROIs, from the first scan for its x-coordinate and rescan for its y-coordinate.

Figure 6 shows the Bland-Altman plots, quantifying the difference between scan and rescan against the average of the measurements. On Bland-Altman plots, bias is detected when the LOA lines do not include zero. Bias is detected in E[*R*_1_] and E[*R*_1_]_bin2_ with this criterion. However, due to the small sample size, the estimated LOA may not accurately represent the true LOA. With larger samples and a Gaussian model, more parameters with detectable bias may emerge.

**Figure 6.**
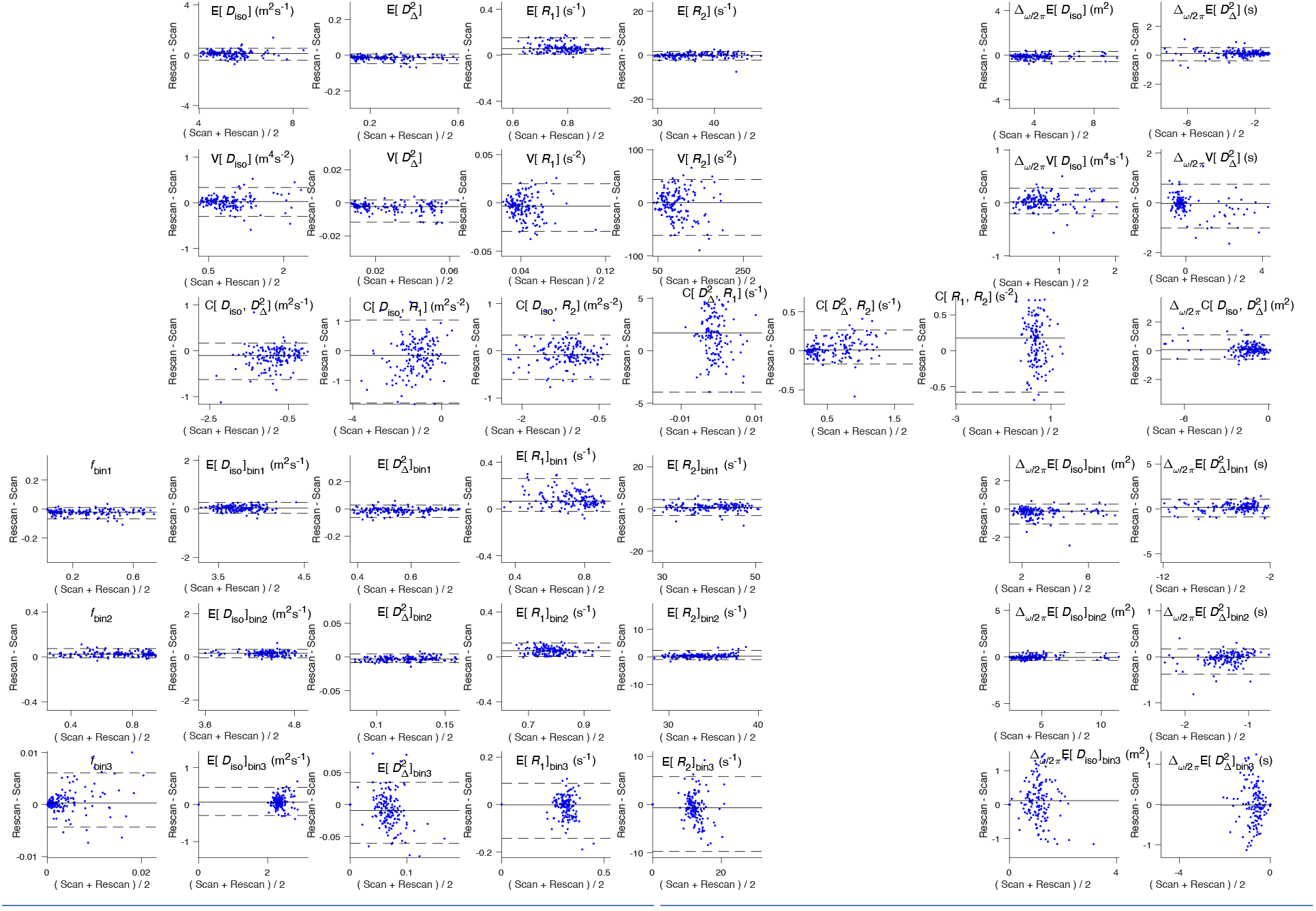
Bland-Altman plots for per-voxel metrics and bin-resolved per-voxel metrics, following the layout of Figure 4. On each plot there is a solid line that depicts the mean “rescan - scan” difference, and two dotted lines depicting the limits of agreement (LOA). Y-axis of plots are scaled to 80% of the maximum average measurement to facilitate comparison, so for poorly performing metrics one or both LOA lines can be out of bounds.

In the correlation plot of E[*R*_1_], we observe significant bias between scan and rescan. We further noticed that, if we color code the time between scan and rescan (not shown), the deviation of the data points from the unity line increases with increasing time between scans, while maintaining a similar slope of one. We identified the source of this bias to be long-term interaction between the sample and the preserving paraformaldehyde solution (detailed in the Discussion). Thus, to isolate the bias related to sample condition from “method-specific” reproducibility, we calculate Pearson’s correlation coefficient for each sample separately, and show the results as box plots (Figure 7). Of interest are metrics with detectable or suspected bias, and we characterize them by their medians and interquartile ranges (IQR). They are E[*R*_1_] (median: 0.972, IQR: 0.961-0.978), E[*R*_1_]_bin1_ (median: 0.975, IQR: 0.937-0.981), E[*R*_1_]_bin2_ (median: 0.946, IQR: 0.937-0.973) and E[*D*_iso_]_bin2_(median: 0.967, IQR: 0.932-0.979). On the distribution of Pearson correlation coefficients, E[*R*_1_] performs similar to E[*D*_iso_] (median: 0.974, IQR: 0.963-0.976), and E[*R*_1_]_bin1_, E[*R*_1_]_bin2_ and E[*D*_iso_]_bin2_ performs similar to E[*R*_2_]_bin2_ (median: 0.966, IQR: 0.949-0.982). E[*D*_iso_] and E[*R*_2_]_bin2_ are both metrics with moderate CCC values (0.938 and 0.943).

**Figure 7.**
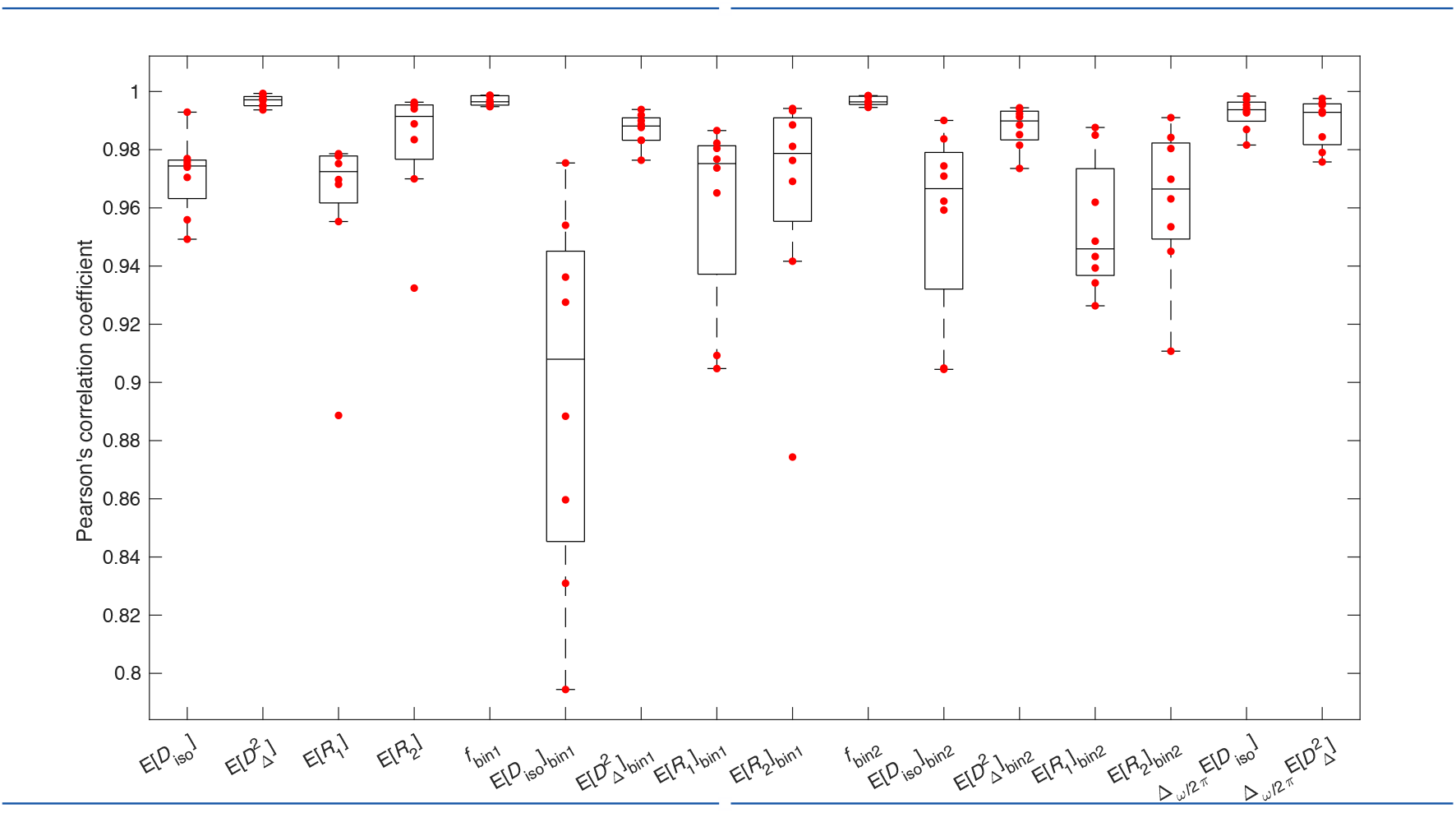
Box plots for per-sample Pearson correlation coefficients of selected metrics.

## 4. Discussion

Diffusion-relaxation correlation MD-MRI enables detailed tissue microstructure characterization by mapping multidimensional distributions of MRI contrasts. This approach enhances sensitivity and specificity by isolating distinct diffusion-relaxation spectral ranges, reducing signal averaging effects. In this study, we investigated the reproducibility of MMD-MRI metrics using ex vivo, preserved mice brains. This study systematically evaluated regional multicomponent diffusion-relaxation properties across white matter tracts, cortical gray matter, and subcortical gray matter, revealing distinct microstructural differences. Additionally, a test-retest analysis demonstrated that the reproducibility of MMD-MRI parameters is comparable to alternative metrics, supporting its reliability for pre-clinical brain imaging.

### 4.1 Regional microstructural features

As seen in previous MMD-MRI studies^28,34,35^, the 2D projections characterize different tissue types well. In WM voxels we find high 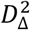 values, and in GM voxels we find low 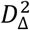 values. More notably, the analysis reveals the presence of multiple distinguishable components within a single voxel, where the variations between these compartments are more subtle than the pronounced differences observed between anisotropic WM and isotropic GM. For example, in the single voxel and ROI-averaged 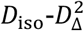 projection of the cortical plate in Figure 3c, it can be seen that the 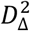 dimension separates the distribution into two. Recently, Yon et al.^35^ also demonstrated a detection of two compartments in GM in in vivo rat brain scans with MMD-MRI 2D projections, separated by *D*_iso_. The reproducibility of these smaller compartments should be investigated in a future study.

### 4.2 Reproducibility comparison with clinical MMD-MRI

The only other scan-rescan study for MMD-MRI was done recently by Manninen et al.^34^ on healthy human participants on a 3T clinical scanner. Reproducibility was quantified using the intraclass correlation coefficient (ICC), and variability was quantified using within-subject coefficient of variation (*CV*_ws_). Due to the small sample size that further separates into two groups (wild type and 5xFAD transgenic mice) of this study, ICC cannot be reliably estimated in the same manner. Using an online calculator^51^ recommended in the literature^52^, for an expected ICC of 0.9, the setup of this study would result in a 95% confidence interval is [0.65, 1.15], which includes three out of the four ICC classes recommended by Koo & Li^53^. We therefore only estimated *CV*_ws_ for our data, separately for the two groups of mouse brains, to facilitate direct comparison with the in vivo MMD-MRI results. A side-by-side comparison of *CV*_ws_ values from Manninen et al. and from the wild type mice in the current study is shown in Figure 8.

**Figure 8.**
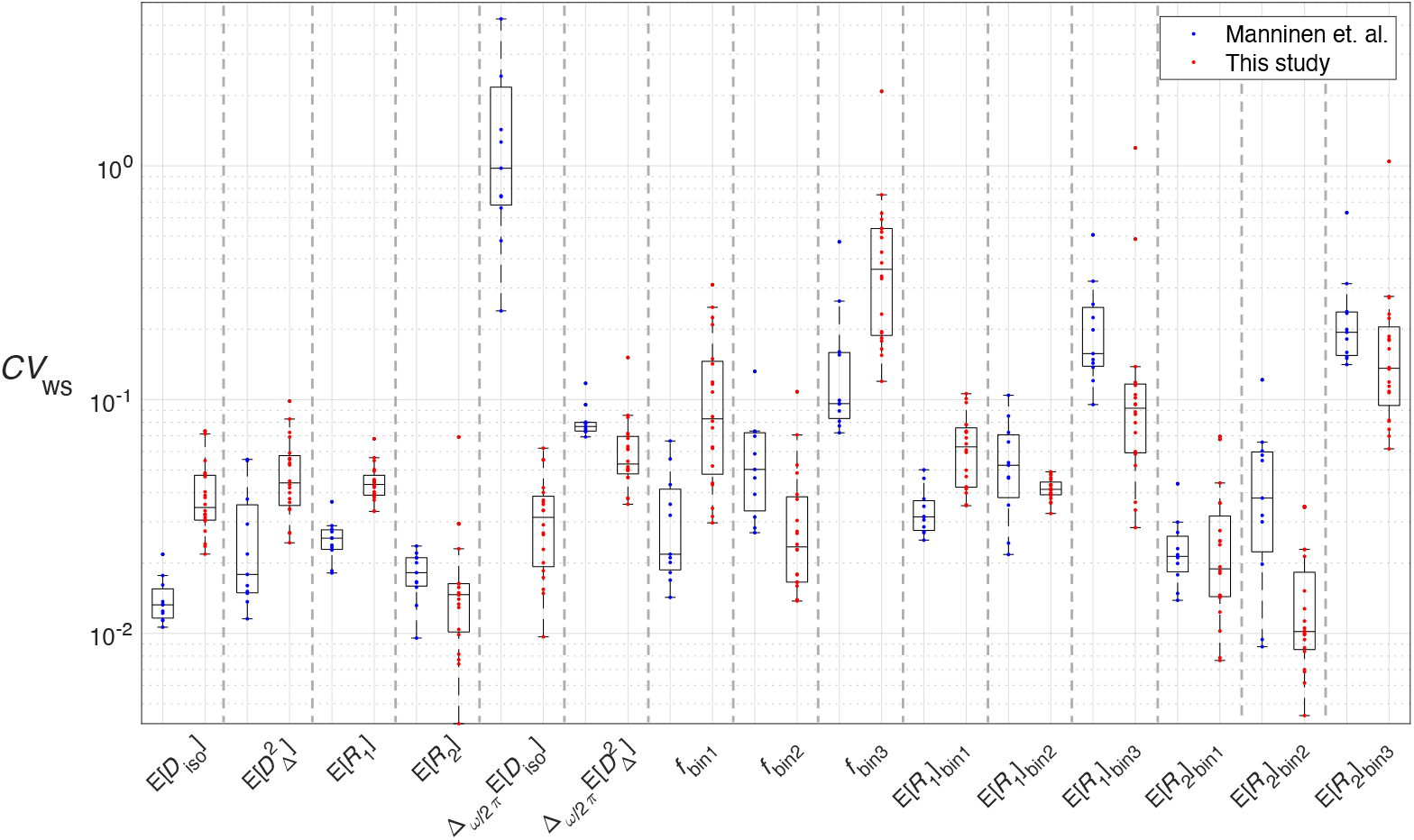
Box plots for ROI-specific *CV*_ws_ for selected MMD-MRI metrics, retrieved from Manninen et al. and calculated for the data in this study. Outlier criterion is value > 1.5x interquartile range from 25^th^ or 75^th^ percentile (box edge).

In general, metrics that performed well in Manninen et al. also performed well in this study. The most striking differences are the much better performance of Δ_*ω/*2*π*_E[*D*_iso_] and the poorer performance of *f*_bin1_ in our study. We attribute the better Δ_*ω/*2*π*_E[*D*_iso_] performance to the larger diffusion encoding frequency range in the acquisition protocol of this study, the *ω*_cent_ of which in Manninen et al. was 6.6-21 Hz and in this study is 36.6-247 Hz. The poorer performance of *f*_bin1_ is likely due to the lower WM content in the mouse brain compared to the human brain, resulting in lower *f*_bin1_ values in this study. In Manninen et al., all ROIs have *f*_bin1_ > 0.3. In our study, for ROIs with *CV*_ws_> 0.1 in the *f*_bin1_ parameter, *f*_bin1_ means are smaller than 0.2. As *CV*_ws_ is a ratio between measurement standard deviation and measurement mean, a smaller denominator naturally gives a larger coefficient of variation.

In terms of reproducibility, although a quantitative comparison cannot be made between the ICCs in that study and the CCCs in this, it is worth noting that in the in vivo study the ICC values of the frequency-sensitive parameters (Δ_*ω/*2*π*_E[*D*_iso_] and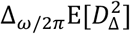) performed the worst among the investigated parameters, while in our study they are among the best. We again attribute the discrepancy to the larger diffusion encoding frequency range in the acquisition protocol of this study.

### 4.3 Reproducibility of variance and covariance metrics

Not investigated in Manninen et al., variance and covariance (V[*x*] and C[*x, y*]) metrics can also be extracted from the bootstrap inversion solution. With the exceptions of V[*D*_*iso*_]and 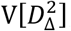,these metrics have lower reproducibility (CCC ≤ 0.9).

In theory, the variances of the solution component distribution describe the diversity in microenvironments in the voxel, but in practice they also describe the uncertainty in the solution space due to the presence of noise^31^. The high reproducibility of V[*D*_*iso*_] and 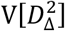 suggests that the MRI acquisition protocol sufficiently samples the diffusion-weighting space and leads to the also high reproducibility of bin fraction estimation (except for bin 3, which contributes insignificantly to the measurement signal in the investigated ROIs). On the contrary, the lower reproducibility of V[*R*_1_] and V[*R*_2_] suggests that the acquisition protocol might be improved by incorporating more variation in repetition time and echo time to stabilize the solution for the relaxation rates.

The covariance metric investigates the relationship between any two MMD-MRI parameters, and the lower reproducibility (0.9 ≥ CCC) suggests a more qualitative interpretation should be taken for these metrics.

Apart from noise sensitivity, partial volume effects also negatively affect the reproducibility of V[*x*] and C[*x, y*] metrics, more than E[*x*] metrics. The exact placement of a voxel on a boundary between different tissue types (e.g. CSF/paraformaldehyde vs GM) affects the ratio of mixing. While the dependence of E[*x*] metrics on mixing ratio is monotonous determined by the two means of the two voxel types, the dependence of V[*x*] and C[*x, y*] metrics is non-linear (parabolic for the case of V[*x*]). For the sake of illustration, consider the *R*_2_ of a voxel with a partial mix of paraformaldehyde and GM (respectively, means: 10 s^-1^ and 30 s^-1^, variances: 10 s^-2^ and 50 s^-2^). While the E[*R*_2_] of the voxel lies between the two means, V[*R*_2_] reaches a maximum of ∼140 s^-2^ at ∼0.5 mixing ratio. The expected absolute difference of E[*R*_2_] and V[*R*_2_] between two scans can then be computed, giving ∼7 s^-1^ and ∼40 s^-2^ respectively.

### 4.5 Systematic bias of MMD-MRI metrics

For any reproducibility study, systematic bias is not usually expected. Here, systematic bias was found for E[*R*_1_] and E[*R*_1_]_bin2_, and by inspection we suspect bias also in 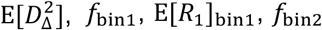 and E[*D*_iso_]_bin2_. It is counterintuitive that some of these metrics can both have high reproducibility and suspected bias at the same time, and we make the argument that it is because the variability in these measurements is low enough such that bias with a magnitude small relative to the scale of the measurement can be detected.

Two possible sources of bias are the change in the flow rate of the vt gas between the first and the second repeat scans (changed from 400 Lph to 200 Lph), and changes in sample condition due to long-time progressive fixation. A follow up experiment scanning one sample twice back-to-back at the two different flow rates was done to disentangle the two factors. Bias was not seen in the new results, leading us to conclude that the cause is chemical degradation. Although the effects of fixation on sample condition time scales relevant to our study (> 4 months after fixation, 3-7 months between scan and rescan) have not been investigated in the literature, relaxation rates have been shown to progressively increase at shorter times^54^, and diffusion rates also have been shown to increase^55^.

### 4.5 Reproducibility comparison with other MRI studies

Coelho et al. investigated the reproducibility of 2D diffusion-*R*_2_ correlation metrics on a human brain in vivo on a clinical scanner^8^. Like MMD-MRI, they used a variable TE and tensor valued encoding imaging sequence. Unlike MMD-MRI’s non-parametric parameter estimation method, they model their signal with the “standard model” of white matter^56^, which in brief is a 3-compartment signal model with diffusion and transverse relaxation contributions, and estimate the model parameters with machine learning inversion. They presented voxelwise CCC values for the model parameters calculated from two repeated scans from a single volunteer. In Table 3, we categorize the standard model parameters into four categories, and do the same for MMD-MRI parameters for a side-by-side comparison of their reproducibility. We make two observations: first, the MMD-MRI parameters in this study perform better, which can be attributed to the ROI-wise evaluation of CCCs, a lengthier and thus a more thorough acquisition protocol, and better gradient hardware. Second, similar to our results, their anisotropy parameter outperforms their diffusivity parameters.

**Table 3.**
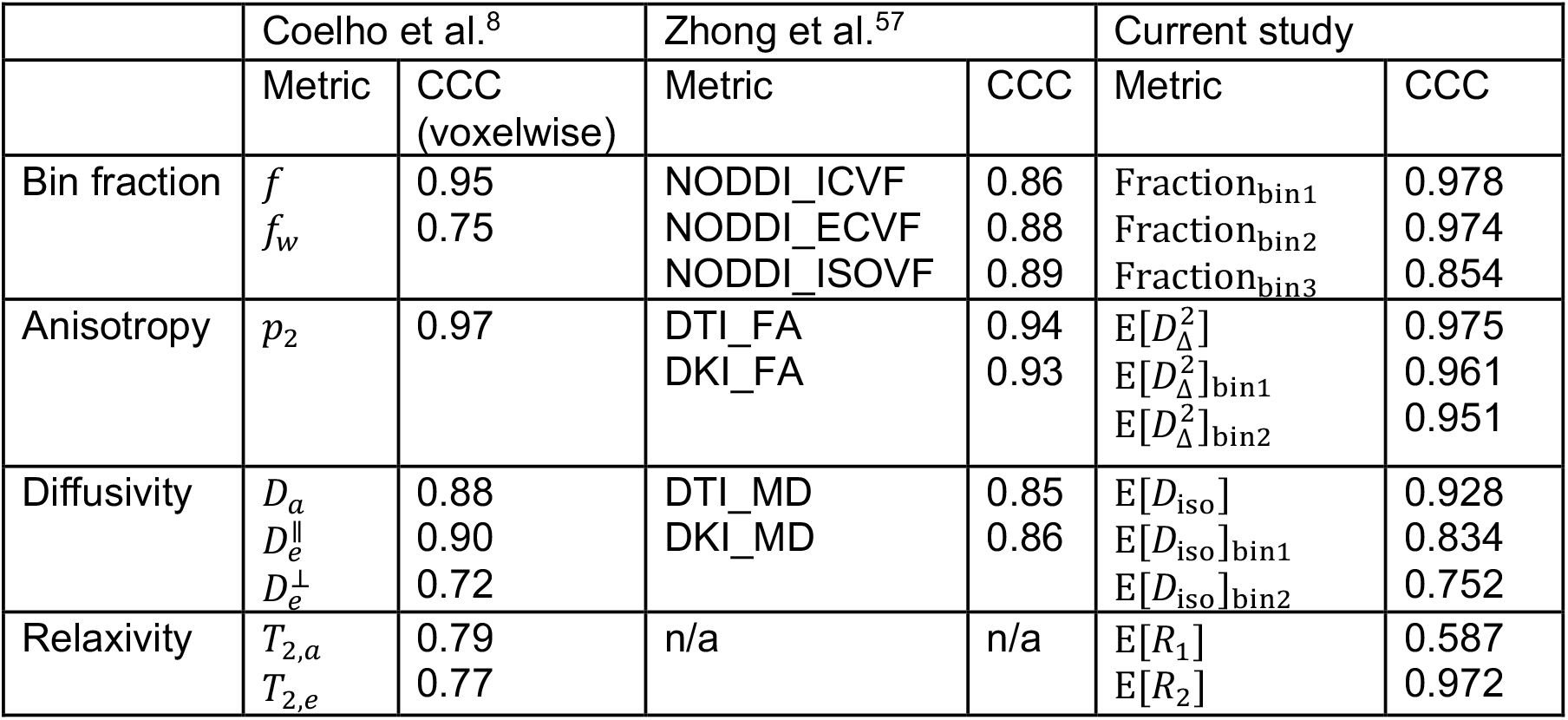
Comparison of CCC reproducibility findings from the current study and two other studies.

Another study by Zhong et al.^57^ investigated scan-rescan reproducibility of diffusion MRI metrics estimated from data acquired with a conventional multi-shell diffusion MRI protocol using four popular diffusion models (DTI^58^, DKI^59^, MAP^60^ and NODDI^6^). Relevant parameters from the four models are also shown in Table 3 for comparison with our results. Again, metrics from our study performed better overall and anisotropy metrics performed better than diffusivity metrics.

### 4.5 Limitations

The small sample size is a limitation that affected the analyses in this study. Aside from prohibiting the use of conventional reproducibility metrics such as ICCs and coefficients of variation, we also opted for a joint analysis of all ROIs altogether instead of individual ROIs separately. It is thus difficult to identify systematic differences, if present, of the reproducibility behavior in individual ROIs.

Despite quality assurance by inspection at every step of the image registration procedure, misregistration unavoidably reduces the reproducibilities calculated in this study. Because we investigate the reproducibility of ROI-averages, the effect for GM regions is expected to be small due to their low surface area-to-volume ratio. On the other hand, WM tract ROIs are generally long and thin. At the MMD-MRI resolution used in this study, mouse WM tracts contain portions that are 1 voxel thin, and thus are more susceptible to misregistration. For the same reason, WM tracts are also susceptible to partial volume effects, a significant portion of which contain a mix of GM and WM.

To minimize these effects, the simplest solution would be to increase the spatial resolution of the MMD-MRI acquisition. However, the resolution of EPI readout is limited by the sample’s apparent transverse relaxation rate 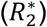 due to its reliance on gradient echoes. Increasing the echo train number to sample a larger k-space, thus increasing spatial resolution, is ineffective, as signal loss occurs at the train’s end. Increasing the number of readout segments could be a workaround, but that would at least double the already lengthy scan time which was impractical for us.

Alternatively, readout methods that are not heavily limited by 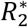 relaxation such as RARE^61^ and GRASE^62^ can be used. However, EPI has the advantage of achieving a shorter echo time, which is important for this study because of the high fast transverse relaxation of the samples.

Outside of MRI methodology solutions, the problem can also be approached from the samples themselves. For example, dissecting the brains into smaller components enables the use of a smaller field of view, which enables larger k-space echo spacing, allowing for faster sampling of *k*-space while keeping the echo train length the same or less. The use of rat brains, which have larger structures than mouse brains, may also be an alternative depending on the scope of the study.

## 5 Conclusion and future work

In this study, we investigated the reproducibility of MMD-MRI metrics on ex vivo mouse brain samples, and showed metrics that are highly repeatable, and metrics that are sensitive to the conditions of the samples measured but otherwise also repeatable when bias is excluded. We offer plausible explanations to the changes in the conditions, which informs future ex vivo MMD-MRI studies, including one we will present in another article on the statistical differences between the two mouse groups used in this study.

Overall, our first order metrics 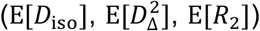,some of the second order metrics 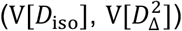, frequency-sensitive metrics 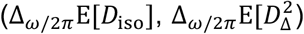, and some of the binned metrics 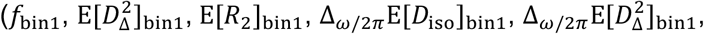 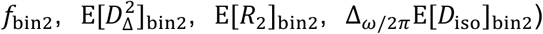 performed similarly or better than the reproducibility of MRI metrics found in the literature.

One of the most significant findings here is the remarkable reproducibility of the frequency-dependent parameters, Δ_*ω/*2*π*_E[*D*_iso_] and 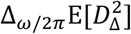.Because of their potential to measure microstructural features through restricted diffusion, methods that probe frequency-dependence have been studied since the 1990s^63^. Yet, the reproducibility of these MMD-MRI metrics has not been demonstrated in a clinical setting.

In fact, as discussed, they have been shown to be unreliable^34^. We hypothesize that the reason those metrics are repeatable here but not on human scanners is the smaller diffusion encoding frequency ranges achievable on typical human scanners (6.6-21 Hz vs 36.6-247 Hz). These results thus motivate further optimization of diffusion encoding waveforms to maximize frequency, and improvement of magnetic gradient hardware^64^ on clinical scanners to enable higher encoding frequencies.

To date, multiple versions of MMD-MRI protocols already exist on clinical scanners^19,33^ with the most recent rendition achieving full brain coverage at 2 mm isotropic resolution on a 3T Philips scanner at a scan time of ∼20 minutes (unpublished). What separates clinical protocols from our preclinical protocol are the limitation of scan time, which restrains the number of parameters that can be sampled, and hardware, which mainly restricts the range of diffusion encoding spectral frequencies that can be probed. These will be points of concerns when attempting to reproduce preclinical results in a clinical setting.

## Acknowledgements

The authors thank Zoltan Tackas for facilitating the acquisition of the data used in this study. This research was partially supported by the Intramural Research Program of the NIH, National Institute on Aging, the Swedish Foundation for Strategic Research (Stiftelsen för Strategisk Forskning; grant no. ITM17-0267), the Swedish Research Council (Vetenskapsrådet; grant nos. 2018-03697, 2022-04422_VR, 21073), and the Research Council of Finland (Project Funding #361370, and Flagship of Advanced Mathematics for Sensing Imaging and Modelling (FAME) #358944).

## Abbreviations

τ_*E*_: echo time
τ_*R*_: repetition time
*ω*: diffusion frequency
b: diffusion encoding tensor
*b*_Δ_: encoding anisotropy
*b*(*ω*): diffusion encoding spectrum
CCC: Lin’s Concordance Correlation Coefficient
ROI: region of interest
LOA: limits of agreement
*CV*_ws_: within-subject coefficient of variation
CSF: cerebrospinal fluid
D: diffusion tensor
*D*_Δ_(*ω*): frequency-dependent normalized diffusion anisotropy
*D*_iso_(*ω*): frequency-dependent isotropic diffusivity
DTD: diffusion tensor distribution
FLASH: fast low-angle shot
GM: gray matter
WM: white matter
MMD: massively multidimensional diffusion
OGSE: oscillating gradient spin echo
*R*_(_: longitudinal relaxation rate
*R*_2_: transversalerelaxation rate
EPI: echo planar imaging;

## ETHICS STATEMENT

The research was conducted according to the principles expressed in the Declaration of Helsinki. The animal study was reviewed and approved by the Animal Committee of the Provincial Government of Southern Finland.

## DATA AND CODE AVAILABILITY

MATLAB source code for preprocessing and Monte-Carlo data inversion is freely available at https://github.com/maximeYon/MMD. The acquisition sequence is available upon reasonable request depending on Paravision versions.

## References

1. Novikov DS. The present and the future of microstructure MRI: From a paradigm shift to normal science. J Neurosci Methods. 2021;351:108947. doi: 10.1016/j.jneumeth.2020.108947

2. Stanisz GJ, Wright GA, Henkelman RM, Szafer A. An analytical model of restricted diffusion in bovine optic nerve. Magn Reson Med. 1997;37(1):103–111. doi: 10.1002/mrm.1910370115

3. Persson F, Söderhjelm P, Halle B. How proteins modify water dynamics. J Chem Phys. 2018;148(21) doi: 10.1063/1.5026861

4. Topgaard D. Multidimensional diffusion MRI. J Magn Reson. 2017;275:98–113. doi: 10.1016/j.jmr.2016.12.007

5. Slator PJ, Palombo M, Miller KL, et al. Combined diffusion-relaxometry microstructure imaging: Current status and future prospects. Magn Reson Med. 2021;86(6):2987–3011. doi: 10.1002/mrm.28963

6. Zhang H, Schneider T, Wheeler-Kingshott CA, Alexander DC. NODDI: Practical in vivo neurite orientation dispersion and density imaging of the human brain. Neuroimage. 2012;61(4):1000–1016. doi: 10.1016/j.neuroimage.2012.03.072

7. Lampinen B, Szczepankiewicz F, Lätt J, et al. Probing brain tissue microstructure with MRI: principles, challenges, and the role of multidimensional diffusion-relaxation encoding. Neuroimage. 2023;282:120338. doi: 10.1016/j.neuroimage.2023.120338

8. Coelho S, Liao Y, Szczepankiewicz F, et al. Assessment of Precision and Accuracy of Brain White Matter Microstructure using Combined Diffusion MRI and Relaxometry. ArXiv. 2025;

9. Stanisz GJ, Henkelman RM. Diffusional anisotropy of T2 components in bovine optic nerve. Magn Reson Med. 1998;40(3):405–410. doi: 10.1002/mrm.1910400310

10. Hürlimann MD, Burcaw L, Song Y-Q. Quantitative characterization of food products by two-dimensional D–T2 and T1–T2 distribution functions in a static gradient. J Colloid Interface Sci. 2006;297(1):303–311. doi: 10.1016/j.jcis.2005.10.047

11. Hürlimann MD, Venkataramanan L. Quantitative Measurement of Two-Dimensional Distribution Functions of Diffusion and Relaxation in Grossly Inhomogeneous Fields. J Magn Reson. 2002;157(1):31–42. doi: 10.1006/jmre.2002.2567

12. Silva MD, Helmer KG, Lee J-H, Han SS, Springer CS, Sotak CH. Deconvolution of Compartmental Water Diffusion Coefficients in Yeast-Cell Suspensions Using Combined T1 and Diffusion Measurements. J Magn Reson. 2002;(1):52–63. doi: 10.1006/jmre.2002.2527

13. Benjamini D, Iacono D, Komlosh ME, Perl DP, Brody DL, Basser PJ. Diffuse axonal injury has a characteristic multidimensional MRI signature in the human brain. Brain. 2021;144(3):800–816. doi: 10.1093/brain/awaa447

14. Kim D, Doyle EK, Wisnowski JL, Kim JH, Haldar JP. Diffusion-relaxation correlation spectroscopic imaging: A multidimensional approach for probing microstructure. Magn Reson Med. 2017;78(6):2236–2249. doi: 10.1002/mrm.26629

15. Benjamini D, Hutchinson EB, Komlosh ME, et al. Direct and specific assessment of axonal injury and spinal cord microenvironments using diffusion correlation imaging. Neuroimage. 2020;221:117195. doi: 10.1016/j.neuroimage.2020.117195

16. Benjamini D, Priemer DS, Perl DP, Brody DL, Basser PJ. Mapping astrogliosis in the individual human brain using multidimensional MRI. Brain. 2022;146(3):1212–1226. doi: 10.1093/brain/awac298

17. Slator PJ, Hutter J, Palombo M, et al. Combined diffusion-relaxometry MRI to identify dysfunction in the human placenta. Magn Reson Med. 2019;82(1):95–106. doi: 10.1002/mrm.27733

18. Naranjo ID, Reymbaut A, Brynolfsson P, et al. Multidimensional Diffusion Magnetic Resonance Imaging for Characterization of Tissue Microstructure in Breast Cancer Patients: A Prospective Pilot Study. Cancers. 2021;13(7):1606.

19. Martin J, Reymbaut A, Schmidt M, et al. Nonparametric D-R1-R2 distribution MRI of the living human brain. Neuroimage. 2021;245:118753. doi: 10.1016/j.neuroimage.2021.118753

20. Callaghan PT, Stepišnik J. Frequency-Domain Analysis of Spin Motion Using Modulated-Gradient NMR. J Magn Reson. 1995;117:118–122. doi: 10.1006/jmra.1995.9959

21. Baron CA, Kate M, Gioia L, et al. Reduction of Diffusion-Weighted Imaging Contrast of Acute Ischemic Stroke at Short Diffusion Times. Stroke. 2015;46(8):2136–2141. doi: 10.1161/STROKEAHA.115.008815

22. Andica C, Hori M, Kamiya K, et al. Spatial Restriction within Intracranial Epidermoid Cysts Observed Using Short Diffusion-time Diffusion-weighted Imaging. Magn Reson Med Sci. 2018;17(3):269–272. doi: 10.2463/mrms.cr.2017-0111

23. Maekawa T, Hori M, Murata K, et al. Differentiation of high-grade and low-grade intra-axial brain tumors by time-dependent diffusion MRI. Magn Reson Imaging. 2020;72:34–41. doi: 10.1016/j.mri.2020.06.018

24. Jespersen SN, Olesen JL, Ianuş A, Shemesh N. Effects of nongaussian diffusion on “isotropic diffusion” measurements: An ex-vivo microimaging and simulation study. J Magn Reson. 2019;300:84–94. doi: 10.1016/j.jmr.2019.01.007

25. Jelescu IO, Veraart J, Fieremans E, Novikov DS. Degeneracy in model parameter estimation for multi-compartmental diffusion in neuronal tissue. NMR Biomed. 2016;29(1):33–47. doi: 10.1002/nbm.3450

26. Kundu S, Barsoum S, Ariza J, et al. Mapping the individual human cortex using multidimensional MRI and unsupervised learning. Brain Commun. 2023;5(6) doi: 10.1093/braincomms/fcad258

27. Lampinen B, Szczepankiewicz F, Mårtensson J, et al. Towards unconstrained compartment modeling in white matter using diffusion-relaxation MRI with tensor-valued diffusion encoding. Magn Reson Med. 2020;84(3):1605–1623. doi: 10.1002/mrm.28216

28. Narvaez O, Svenningsson L, Yon M, Sierra A, Topgaard D. Massively Multidimensional Diffusion-Relaxation Correlation MRI. Frontiers in Physics. 2022;9 doi: 10.3389/fphy.2021.793966

29. Topgaard D. Multiple dimensions for random walks. J Magn Reson. 2019;306:150–154. doi: 10.1016/j.jmr.2019.07.024

30. Narvaez O, Yon M, Jiang H, et al. Nonparametric distributions of tensor-valued Lorentzian diffusion spectra for model-free data inversion in multidimensional diffusion MRI. J Chem Phys. 2024;161(8) doi: 10.1063/5.0213252

31. Prange M, Song Y-Q. Quantifying uncertainty in NMR T2 spectra using Monte Carlo inversion. J Magn Reson. 2009;196(1):54–60. doi: 10.1016/j.jmr.2008.10.008

32. de Almeida Martins JP, Topgaard D. Multidimensional correlation of nuclear relaxation rates and diffusion tensors for model-free investigations of heterogeneous anisotropic porous materials. Sci Rep. 2018;8(1):2488. doi: 10.1038/s41598-018-19826-9

33. Johnson JTE, Irfanoglu MO, Manninen E, et al. In vivo disentanglement of diffusion frequency-dependence, tensor shape, and relaxation using multidimensional MRI. Hum Brain Mapp. 2024;45(7):e26697. doi: 10.1002/hbm.26697

34. Manninen E, Bao S, Landman BA, Yang Y, Topgaard D, Benjamini D. Variability of multidimensional diffusion–relaxation MRI estimates in the human brain. Imaging Neuroscience. 2024;2:1–24. doi: 10.1162/imag_a_00387

35. Yon M, Narvaez O, Topgaard D, Sierra A. In vivo rat-brain mapping of multiple gray matter water populations using nonparametric D(ω)-R1-R2 distributions MRI. NMR Biomed. 2025;38(1):e5286. doi: 10.1002/nbm.5286

36. Oakley H, Cole SL, Logan S, et al. Intraneuronal β-Amyloid Aggregates, Neurodegeneration, and Neuron Loss in Transgenic Mice with Five Familial Alzheimer’s Disease Mutations: Potential Factors in Amyloid Plaque Formation. J Neurosci. 2006;26(40):10129–10140. doi: 10.1523/jneurosci.1202-06.2006

37. Yon M, Narvaez O, Topgaard D, Sierra A. In vivo rat brain mapping of multiple gray matter water populations using nonparametric D(ω)-R1-R2 distributions MRI. NMR Biomed. 2025;38(1):e5286. doi: 10.1002/nbm.5286

38. Jiang H, Svenningsson L, Topgaard D. Multidimensional encoding of restricted and anisotropic diffusion by double rotation of the q vector. Magn Reson. 2023;4(1):73–85. doi: 10.5194/mr-4-73-2023

39. Veraart J, Novikov DS, Christiaens D, Ades-aron B, Sijbers J, Fieremans E. Denoising of diffusion MRI using random matrix theory. Neuroimage. 2016;142:394–406. doi: 10.1016/j.neuroimage.2016.08.016

40. Tournier JD, Smith R, Raffelt D, et al. MRtrix3: A fast, flexible and open software framework for medical image processing and visualisation. Neuroimage. 2019;202:116137. doi: 10.1016/j.neuroimage.2019.116137

41. Smith SM, Jenkinson M, Woolrich MW, et al. Advances in functional and structural MR image analysis and implementation as FSL. Neuroimage. 2004;23:S208–S219. doi: 10.1016/j.neuroimage.2004.07.051

42. Nilsson M, Szczepankiewicz F, Lampinen B. An open-source framework for analysis of multidimensional diffusion MRI data implemented in MATLAB. Proc Int Soc Magn Reson Med Sci Meet Exhib. Accessed Jan 26, 2024,

43. Lundell H, Nilsson M, Dyrby TB, et al. Multidimensional diffusion MRI with spectrally modulated gradients reveals unprecedented microstructural detail. Sci Rep. 2019;9(1):9026. doi: 10.1038/s41598-019-45235-7

44. Topgaard D. Diffusion tensor distribution imaging. NMR Biomed. 2019;32(5):e4066. doi: 10.1002/nbm.4066

45. Ellmore TM, Murphy SM, Cruz K, Castriotta RJ, Schiess MC. Averaging of diffusion tensor imaging direction-encoded color maps for localizing substantia nigra. Computers in Biology and Medicine. 2014;51:104–110. doi: 10.1016/j.compbiomed.2014.05.004

46. Avants BB, Tustison NJ, Song G, Cook PA, Klein A, Gee JC. A reproducible evaluation of ANTs similarity metric performance in brain image registration. Neuroimage. 2011;54(3):2033–2044. doi: 10.1016/j.neuroimage.2010.09.025

47. Lawrence IKL. A Concordance Correlation Coefficient to Evaluate Reproducibility. Biometrics. 1989;45(1):255–268. doi: 10.2307/2532051

48. McBride GB. A Proposal for Strength-of-agreement Criteria for Lin’s Concordance Correlation Coefficient. NIWA Client Report: HAM2005-062. 2005;

49. Klein R. Bland-Altman and Correlation Plot (https://www.mathworks.com/matlabcentral/fileexchange/45049-bland-altman-and-correlation-plot). xAccessed January 25, 2025.

50. Gerke O. Nonparametric Limits of Agreement in Method Comparison Studies: A Simulation Study on Extreme Quantile Estimation. Int J Environ Res Public Health. 2020;17(22):8330.

51. Arifin W. Sample size calculator (web). Accessed Feb 03, 2025. https://wnarifin.github.io/ssc/ssicc.html

52. Monti CB, Ambrogi F, Sardanelli F. Sample size calculation for data reliability and diagnostic performance: a go-to review. Eur Radiol Exp. 2024;8(1):79. doi: 10.1186/s41747-024-00474-w

53. Koo TK, Li MY. A Guideline of Selecting and Reporting Intraclass Correlation Coefficients for Reliability Research. J Chiropr Med. 2016;15(2):155–163. doi: 10.1016/j.jcm.2016.02.012

54. Shatil AS, Uddin MN, Matsuda KM, Figley CR. Quantitative Ex Vivo MRI Changes due to Progressive Formalin Fixation in Whole Human Brain Specimens: Longitudinal Characterization of Diffusion, Relaxometry, and Myelin Water Fraction Measurements at 3T. Front Med. 2018;5 doi: 10.3389/fmed.2018.00031

55. Shepherd TM, Thelwall PE, Stanisz GJ, Blackband SJ. Aldehyde fixative solutions alter the water relaxation and diffusion properties of nervous tissue. Magn Reson Med. 2009;62(1):26–34. doi: 10.1002/mrm.21977

56. Novikov DS, Fieremans E, Jespersen SN, Kiselev VG. Quantifying brain microstructure with diffusion MRI: Theory and parameter estimation. NMR Biomed. 2019;32(4):e3998. doi: 10.1002/nbm.3998

57. Zhong J, Liu X, Hu Y, et al. Robustness of Quantitative Diffusion Metrics from Four Models: A Prospective Study on the Influence of Scan-Rescans, Voxel Size, Coils, and Observers. J Magn Reson Imaging. 2024;60(4):1470–1483. doi: 10.1002/jmri.29192

58. Basser PJ, Mattiello J, LeBihan D. MR Diffusion Tensor Spectroscopy and Imaging. Biophys J. 1994;66:259–267.

59. Jensen JH, Helpern JA, Ramani A, Lu H, Kaczynski K. Diffusional kurtosis imaging: The quantification of non-gaussian water diffusion by means of magnetic resonance imaging. Magn Reson Med. 2005;53(6):1432–1440. doi: 10.1002/mrm.20508

60. Özarslan E, Koay CG, Shepherd TM, et al. Mean apparent propagator (MAP) MRI: A novel diffusion imaging method for mapping tissue microstructure. Neuroimage. 2013;78:16–32. doi: 10.1016/j.neuroimage.2013.04.016

61. Hennig J, Nauerth A, Friedburg H. RARE imaging: A fast imaging method for clinical MR. Magn Reson Med. 1986;3(6):823–833. doi: 10.1002/mrm.1910030602

62. Feinberg DA, Oshio K. GRASE (gradient- and spin-echo) MR imaging: a new fast clinical imaging technique. Radiology. 1991;181(2):597–602. doi: 10.1148/radiology.181.2.1924811

63. Stepišnik J. Time-dependent self-diffusion by NMR spin-echo. Physica B Condens Matter. 1993;183(4):343–350. doi: 10.1016/0921-4526(93)90124-O

64. Vachha B, Huang SY. MRI with ultrahigh field strength and high-performance gradients: challenges and opportunities for clinical neuroimaging at 7 T and beyond. Eur Radiol Exp. 2021;5(1):35. doi: 10.1186/s41747-021-00216-2

